# Rapid acting antidepressant (2R,6R)-hydroxynorketamine (HNK) targets glucocorticoid receptor signaling: a longitudinal cerebrospinal fluid proteome study

**DOI:** 10.1101/2020.09.03.280834

**Authors:** David P. Herzog, Natarajan Perumal, Caroline Manicam, Giulia Treccani, Jens Nadig, Milena Rossmanith, Jan Engelmann, Tanja Jene, Annika Hasch, Michael A. van der Kooij, Klaus Lieb, Nils C. Gassen, Franz H. Grus, Marianne B. Müller

**Author notes:** Shared first authorship in alphabetical order (David P. Herzog, Natarajan Perumal). David P. Herzog, MD, Laboratory of Translational Psychiatry, Department of Psychiatry and Psychotherapy, Johannes Gutenberg University Medical Center Mainz, Hanns-Dieter-Hüsch-Weg 19, 55128 Mainz, Germany. **Conflict of interest statement:** The authors have declared that no conflict of interest exists.

## Abstract

Delayed onset of antidepressant action is a shortcoming in depression treatment. Ketamine and its metabolite (2R,6R)-hydroxynorketamine (HNK) have emerged as promising rapid-acting antidepressants. However, their mechanism of action remains unknown. In this study, we first described the anxious and depression-prone inbred mouse strain, DBA/2J, as a animal model to assess the antidepressant-like effects of ketamine and HNK *in vivo.* To decode the molecular mechanisms mediating HNK’s rapid antidepressant effects, a longitudinal cerebrospinal fluid (CSF) proteome profiling of its acute and sustained effects was conducted using an unbiased, hypothesis-free mass spectrometry-based proteomics approach. A total of 387 proteins were identified, with a major implication of significantly differentially expressed proteins in the glucocorticoid receptor (GR) signaling pathway, providing evidence for a link between HNK and regulation of the stress hormone system. Mechanistically, we identified HNK to repress GR-mediated transcription and reduce hormonal sensitivity of GR *in vitro.* In addition, mammalian target of rapamycin (mTOR) and brain-derived neurotrophic factor (BDNF) were predicted to be important upstream regulators of HNK treatment. Our results contribute to precise understanding of the temporal dynamics and molecular targets underlying HNK’s rapid antidepressant-like effects, which can be used as a benchmark for improved treatment strategies for depression in future.

## Introduction

A well-recognized shortcoming of currently available pharmacological agents in the treatment of major depressive disorder (MDD) is that significant difference from placebo can only be reliably identified 4-6 weeks after treatment onset (1). This therapeutic delay of classical antidepressants is potentially dangerous since suicide risk is the highest in the early treatment period when the disease severity is at its greatest (2). The absence of rapid positive feedback also encourages non-compliance, thus increasing duration of the depressive episode and the overall costs for treating this disease (3). In addition, the lack of biomarkers for disease monitoring or drug choice in the clinical setting is a major drawback in depression treatment (4).

In search for putative targets mediating a more rapid antidepressant response, *N*-methyl-D-aspartate (NMDA) receptor antagonists, with ketamine as a prototypical agent, have gained increasing interest during the last years (5). Sub-anesthetic doses of the narcotic agent ketamine have been shown to produce both rapid (within hours) and sustained (up to 1 week) antidepressant effects in patients (6) and rodents (7). In addition, there is convincing evidence from clinical trials that ketamine is also effective in patients with treatment-resistant depression (8). Besides its activity at the NMDA receptor, several putative molecular mechanisms through which the antidepressant-like effects of ketamine are mediated have been discovered, including the involvement of mechanistic target of rapamycin complex 1 (mTORC1) signaling and acute modulation of brain-derived neurotrophic factor (BDNF) release (9).

Despite its proven antidepressant efficacy even in difficult-to-treat patients, a broad range of side effects, including strong sedation, dissociation, nausea, and the risk of addiction have prevented ketamine to be widely used so far (10). Recent preclinical studies indicate that the ketamine metabolite, (2*R*,6*R*)-hydroxynorketamine (HNK), retains the rapid and sustained antidepressant-like effects in rodents, but lacks its dissociative-like properties and abuse potential (11-13). Therefore, HNK is a potential prototypical candidate to be further tested and developed as a rapid-acting antidepressant agent. However, for HNK, which does not block NMDA receptors like ketamine (14), the molecular signaling mechanisms still remain largely unknown (15). In addition, the antidepressant-like effects of HNK could not always be reproduced by research groups around the world. We summarized the evidence in a recent review (16). To date, systematic studies aiming at the hypothesis-free identification of proteins and signaling pathways involved in HNK’s rapid onset of action are yet to be reported.

Despite considerable efforts, biomarker candidates to support clinical diagnosis or treatment response that could finally support decision-making within MDD diagnosis and therapy guidelines are still lacking (17). Access to the organ of interest as source for biomaterial, i.e. the brain, is restricted, if not impossible. Data on peripheral blood-based biomarkers and molecular candidates have been relatively inconclusive so far, and peripheral changes are remarkably different from the molecular changes that are observed within the central nervous system (e.g. (18)). Cerebrospinal fluid (CSF) is an accessible biological fluid in both humans and rodents that is circulating in close vicinity to the brain. In the past decade, technological advancement and the increasing quality of unbiased “omics” approaches have facilitated the use of CSF for biomarker research and investigations into disease-related pathophysiological processes. However, in contrast to other mental diseases such as neurodegenerative disorders where CSF is traditionally an established source for biomarker research, only very few investigations have used CSF to monitor the changes in the context of antidepressant treatment.

In 2014, Onaivi *et al.* described an animal experimental approach using intraventricular cannulation to enable serial CSF withdrawal in conscious mice (19). The suitability of this approach was validated by revealing specific proteomic signatures in murine CSF samples following acute stress and acute cocaine administration (19). In the current study, we adopted this method to focus on temporal changes of the CSF proteome induced by systemic administration of an antidepressant-like dosage of HNK. As animal model system, we chose male DBA/2J mice. This inbred mouse strain is characterized by high innate anxiety, which can be successfully treated by commonly used antidepressants (18, 20, 21). Moreover, the DBA/2J mice display a reduced inhibitory HPA axis feedback (22) closely mimicking this impairment in depressed humans. Choosing an antidepressant-responsive strain enables experiments to be performed under baseline conditions i.e. without the need to apply additional and potentially confounding stressors to shift the behavioral phenotype of the animals to a depressive-like state to model disease-like conditions (18). Here, we performed serial CSF sample collection from conscious and freely moving mice in a longitudinal study design to identify the proteome changes closely associated with changes in brains states following administration of HNK over time. Changes in the CSF proteome and differential expression of proteins associated with the acute (4 hours after injection) and sustained (1 week after injection) antidepressant-like effects of a single injection of HNK or saline were further investigated employing a hypothesis-free, unbiased mass spectrometry-based proteomics approach.

## Results

### Behavioral profiling of ketamine and HNK in DBA/2J mice

With this behavioral experiment, we wanted to first validate that DBA/2J inbred mice are a suitable strain and model organism to investigate the antidepressant-like effects of ketamine and HNK. DBA/2J mice displayed a depression-like phenotype under control conditions with high immobility scores in the FST (Figure 1A and B at 4h and 1w, respectively; saline-treated animals). At both *acute* and *sustained* time points, ketamine and HNK induced significant antidepressant-like effects by reducing immobility time (Figure 1A and B). We detected no statistically significant differences between the effects of HNK and ketamine at both time points (Figure 1A and B).

**Figure 1.**
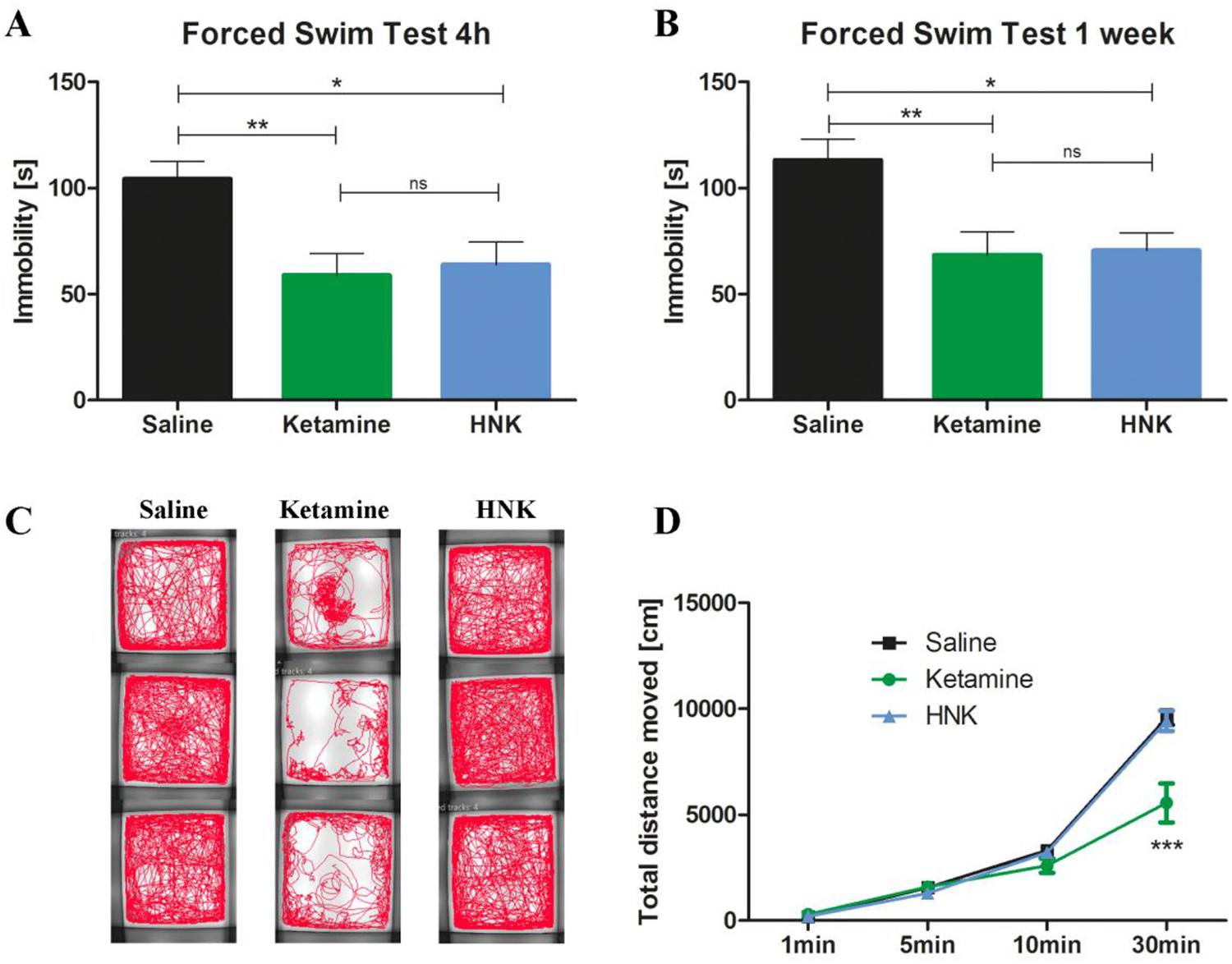
Antidepressant-like effects and side-effects of ketamine and HNK. (A) 4 hours after injection, ketamine [t=3.32, p<0.01] and HNK [t=2.96, p<0.05] reduced immobility time in the FST [F=7.44, p=0.0019, n=10 per group]. (B) 1 week after injection, ketamine [t=3.24, p<0.01] and HNK [t=3.08, p<0.05] reduced immobility time in the FST [F=6.67, p=0.0044, n = 10 per group]. (C+D) Ketamine affected locomotor activity [n=10 per group]. (C) Ketamine-treated mice show an altered tracking profile [3 examples per group]. (D) Ketamine decreased locomotor activity [Interaction: F=13.32, p<0.001; time: F=388.8, p<0.001; treatment: F=7.65, p<0.01] compared to HNK [t=7.79, p<0.001] and saline [t=8.10, p<0.001]. ns not statistically significant, HNK (2R,6R)-hydroxynorketamine, * p<0.05, ** p<0.01, *** p<0.001, bars are depicted as mean ± SEM, one-way ANOVA with Bonferroni posttest in A+B, two-way repeated-measures ANOVA with Bonferroni posttest in D.

Locomotor activity is known to be impaired by ketamine treatment and constitutes an important side-effect of ketamine (10). Indeed, ketamine-induced reduction in general locomotor activity could be visualized by an altered tracking profile in the OF test during 30 minutes of post-injection recording (Figure 1C). Furthermore, ketamine reduced the total distance travelled, compared to HNK and saline injection (Figure 1D).

### Acute and sustained effects of HNK on CSF proteome

CSF is in close vicinity to the brain, and there is a continuous exchange and molecular crosstalk between brain tissue and CSF. Because of the clear behavioral effects of HNK and the fact that so far, data about its mechanism of action remain fragmentary, we decided to exclusively focus on HNK and analyze the CSF samples obtained from conscious DBA/2J mice treated with HNK or saline. As part of the method validation, we investigated whether the CSF withdrawal procedure was stressful for the animal by measurement of plasma CORT concentrations in an independent experiment. The mean peak level of CORT from mice which underwent the CSF withdrawal procedure was 84.84 ng/ml (Figure S1), not exceeding average plasma CORT concentrations reached following exposure to a novel environment, such as a commonly used OF test (23).

In the main experiment, CSF samples were pooled (Supplemental data A), processed, and subjected to first dimensional gel electrophoresis (1DE), as depicted in Figure S2. In total, 387 proteins were identified from both *Mus musculus* (309 proteins) and *Homo sapiens* (229 proteins) databases by label-free quantification at a false discovery rate (FDR) of 1% (Figure 2A; full data in Supplemental data C-E). As many as 151 proteins were found to be overlapping in both databases, as shown in the Venn diagram (Figure 2A). The use of both mouse and human UniProt databases maximized protein identification due to limited annotations in the mouse database (17 001 proteins) compared to the human database (20 410 proteins).

**Figure 2.**
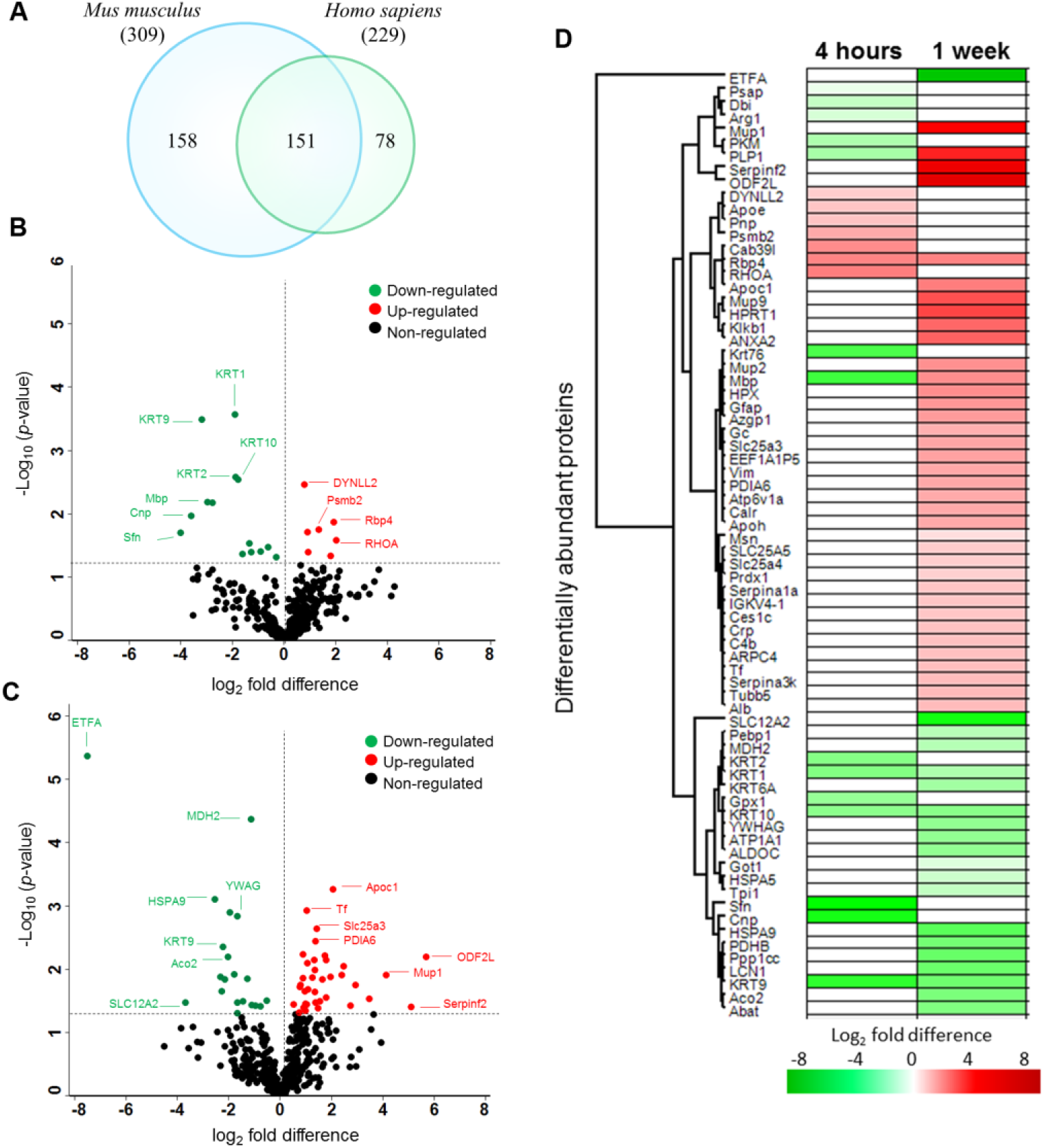
Acute and sustained effects of HNK on CSF proteome. (A) Venn diagram depicts the total number of proteins identified in both mouse and human databases in the CSF samples. Volcano plot illustrates significantly differentially abundant CSF proteins after (B) 4 hours and (C) 1 week HNK administration compared to saline. The negative log10 (p-value) is plotted against the log2 (fold change: HNK/ saline). The non-axial horizontal line denotes p = 0.05, which is our significance threshold (prior to logarithmic transformation). Significantly upregulated proteins are plotted in red, while proteins that were downregulated by HNK are represented in green. (D) The hierarchical clustering of the differentially expressed CSF proteins at 4 hours and 1 week after HNK treatment displayed in a heat map. The upregulated proteins are shown in red and the downregulated proteins are in green, with log2 fold change regulation of intensity displayed in color (from -8 (green) to 8 (red).

Among these murine CSF proteins, 25 and 65 proteins were found to be differentially expressed at 4 hours (Figure 2B) and 1 week (Figure 2C) after HNK treatment compared to saline (for comprehensive list see Supplemental data F and Figure S3). Nine proteins were upregulated following 4 hours HNK administration namely cytoskeletal structural protein dynein light chain 2, cytoplasmatic (Dynll2), proteasome subunit beta type 2 (Psmb2), apolipoprotein (Apoe), and spine growth modulator protein transforming protein RhoA (Rhoa). Sixteen proteins were downregulated following 4 hours HNK versus saline injection (Figure 2C), mainly cytoskeletal keratins consisting of keratin type II cytoskeletal 1, 2 peridermal, 2 oral, 9, 10 (KRT1, KRT2, KRT76, KRT9, KRT10), myelin proteins composed of myelin basic protein (Mbp) and myelin proteolipid protein (Plp1), signaling protein 14-3-3 protein sigma (Sfn), and myelinotrophic and neurotrophic factor prosaposin (Psap). One week following HNK injection, we found 41 proteins to be upregulated (Figure 2C). Examples of these proteins were Mbp, Plp1, glial fibrillary acidic protein (Gfap), amino acid transporters ADP/ATP translocase 2 (SLC25A5) and 1 (Slc25a4), cytoskeletal proteins actin-related protein 2/3 complex subunit 4 (Arpc4), vimentin (Vim), tubulin beta-5 chain (Tubb5), plasma kallikrein (Klkb1), moesin (Msn), and transcription factor putative elongation factor 1-alpha-like 3 (EEF1A1P5). A total of 24 proteins were downregulated comprising mainly cytoskeletal keratins (KRT1, 9, 6A, 10), mitochondrial proteins electron transfer flavoprotein subunit alpha (ETFA), malate dehydrogenase (Mdh2), stress-70 protein (HSPA9), 4- aminobutyrate aminotransferase (Abat), aconitate hydratase (Aco2), pyruvate dehydrogenase E1 component subunit beta (PDHB), Na/K-transporting ATPase subunit alpha-1 (ATP1A1), cholinergic neuron ATP-binding protein in neurons phosphatidylethanolamine-binding protein 1 (Pebp1), and chaperone heat shock protein a5 (HSPA5). Further clustering of these differentially expressed CSF proteins depicted as heat maps with unsupervised hierarchical clustering demonstrated the segregation of identified proteins into two major clusters according to the respective time points (Figure 2D). Three distinct protein expression profiles were observed. First, the expression of a cluster pf proteins remained the same at both 4h and 1w composed of down-regulation of KRT1, KRT9, and KRT10, and the up-regulation of Rbp4. The second cluster consisted of proteins that were upregulated following 1w of HNK administration comprising Mbp and PLP1. The third cluster was characterized by expression of proteins that were only regulated exclusively at either time point. Conducting a protein-protein interaction network (PPI) analysis, we could show the localization in particular cellular compartments and molecular types (Figure S3). Interestingly, the protein group observed to have the highest interaction parterns consisted of keratins (Figure S3).

In an attempt to further explore the functional relevance of the proteins identified to be differentially expressed, canonical pathway enrichment was determined employing the IPA tool using a cut-off of p<0.05. The 21 proteins following 4 hours HNK injection were significantly associated with the glucocorticoid receptor (GR) signaling (*p* = 1.52E-03; Figure 3A). On the other hand, 9 top canonical pathways were found to be differentially regulated by the 57 differentially expressed proteins following 1 week HNK treatment, as shown in Figure 3A. Among these, the acute phase response signaling (*p* = 5.30E-11) was the most significantly implicated pathway, followed by unfolded protein response (*p* = 3.07E-04), clathrin-mediated endocytosis signaling (*p* = 2.17E-03), GR signaling (*p* = 2.35E-03), 14-3-3 mediated signaling (*p* = 3.68E-03), glycolysis I (*p* = 1.21E-02), gluconeogenesis (*p* = 1.21E-02), and signaling by the Rho family GTPase (*p* = 2.13E-02). Noteworthy, the GR signaling-associated proteins at both time points comprised mainly a cluster of keratins and heat shock proteins. Among the keratins, KRT1, KRT9, and KRT 10 were consistently downregulated at both time points, whereas KRT76 and KRT2 were exclusively downregulated at 4 hours and KRT6A at 1 week after HNK treatment (Figure 3B; Supplemental data G).

**Figure 3.**
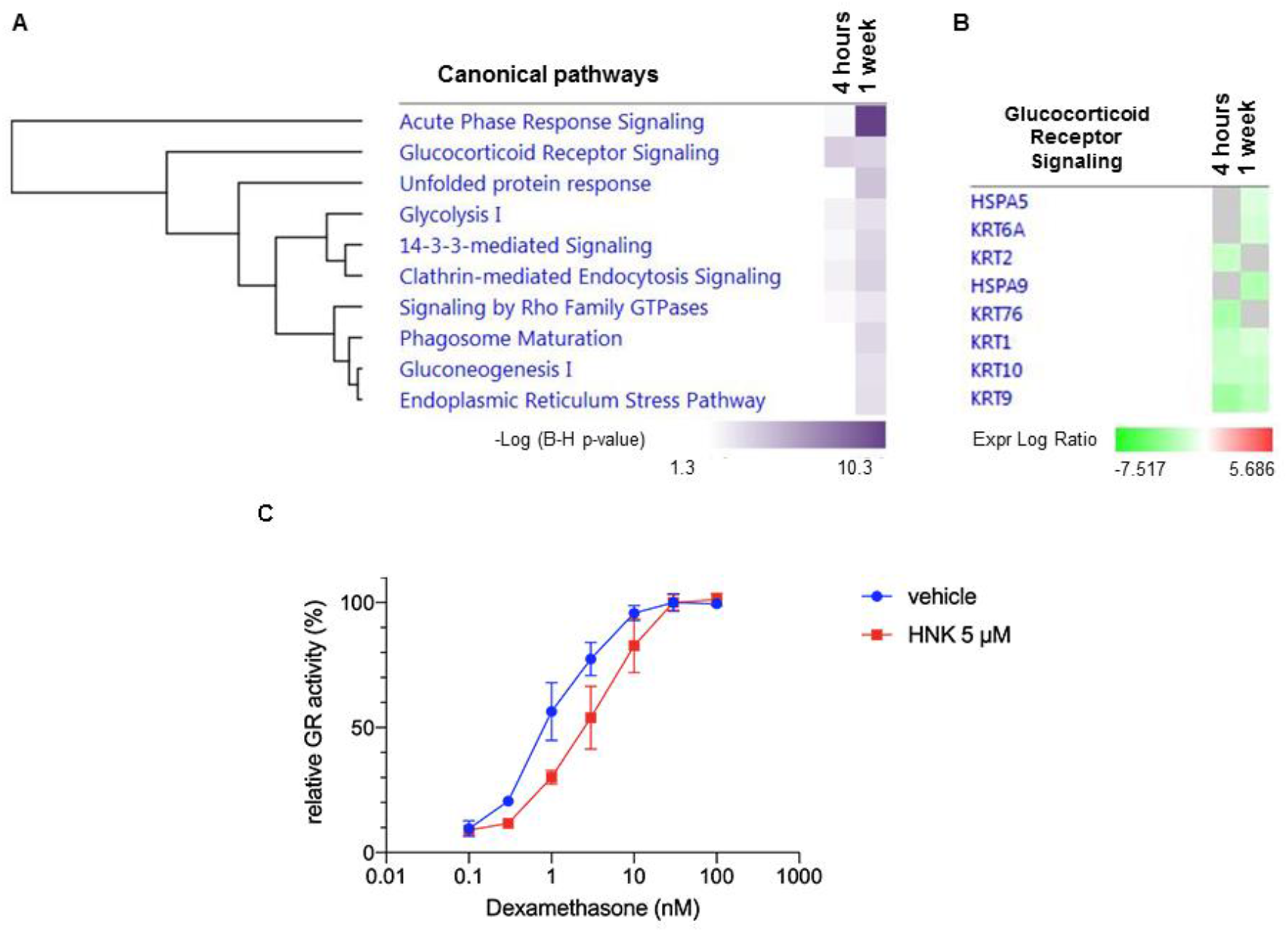
Top enriched canonical pathways of the differentially expressed proteins significantly affected by HNK treatment. (A) The significantly enriched canonical pathways determined by IPA’s default threshold [-log (p-value) >1.3] between the differentially expressed proteins identified in our datasets and the molecules in the respective pathways at 4h and 1w time-points following HNK treatment. (B) Proteins associated with glucocorticoid receptor signaling pathway and their expression profiles at 4 h and 1 w. The intensity of the color (green) indicates the degree of downregulation of each protein. (C) Relative GR activity was reduced through HNK (5 μM) treatment compared to vehicle control administration (p < 0.05 for 1.0 nM dexamethasone treatment). Relative receptor activity represents firefly luciferase activity (MMTV-promoter) normalized to Gaussia luciferase activity and is represented to the activity at saturating 30 nM dexamethasone. 2way ANOVA followed by Bonferroni correction was applied.

### HNK represses GR-mediated transcription and reduces hormonal sensitivity of GR *in vitro*

We were interested how HNK affected GR translational activity and GR response to glucocorticoids in neuronal cells. We performed an *in vitro* gene reporter assay assessing GRE-driven transcriptional activity by transient transfection of MMTV-Luciferase construct into HT-22 mouse hippocampal neuronal cells and found that HNK represses GR-mediated transcription in a glucocorticoid (DEX) dependent manner (Figure S4). We detected no direct effect of HNK on GR-activity, when steroids were withdrawn from culture medium (Figure S4). Moreover, we found a reduced hormonal sensitivity of GR when cells were co-treated with HNK compared to control condition (Figure 3C).

Figure 4 depicts the further analysis of the top disease and biological functions associated with the differentially expressed CSF proteins at both time points. At 4h after HNK treatment, myelination (*p* = 2.43E-03; Figure 4B), regeneration of neurons (*p* = 4.32E-03), and organization of filaments (*p* = 1.04E-02) were found to be significantly affected. The subacute effect of HNK administration after 1w demonstrated that the proteins found to be differentially expressed were significantly associated with top eight networks involved in Alzheimer disease (*p* = 1.93E-07; Figure 4C), organization of filaments (*p* = 6.36E-05; Figure 4D), schizophrenia (*p* = 2.36E-04), gluconeogenesis (*p* = 7.17E-04), neurological signs (*p* = 7.35E-04), proliferation of neuroglia (*p* = 2.01E-03), Parkinson's disease (*p* = 4.03E-03), and regeneration of neurons (*p* = 1.75E-02). Finally, we analyzed the predicted upstream regulators to identify the molecules, including transcription factors, which were significantly involved in the regulation of the differentially expressed proteins in both groups. Interestingly, at the 4h time point, the major regulators were all inhibited comprising insulin like growth factor 1 (IGF1) (*p* = 7.7E-05; z-score = -2.18), mTOR (*p* = 1.2E-05; z-score = - 0.475), tumor necrosis factor (TNF) (*p* = 1.6E-04; z-score = -0.743) and transforming growth factor beta 1 (TGFβ1) (*p* = 2.5E-02; z-score = -0.152), as depicted in Figure 5A. However, at the 1 w time point following HNK treatment, five upstream regulators were predicted to be activated, which were interleukin 6 (IL6) (*p* = 1.2E-05; z-score = 1.337), IGF1 (*p* = 1.7E-04; z-score = 1.28), TNF (*p* = 3.9E-03; z-score = 0.742), mTOR (*p* = 1.2E-03; z-score = 0.243), and BDNF (*p* = 1.2E-04; z-score = 0.128; Figure 5B). Three regulators comprising transforming growth factor alpha (TGFα) (*p* = 5.0E-05; z-score = -0.257) and TGFβ1 (*p* = 3.4E-04; z-score = -0.24), and androgen receptor (AR) (*p* = 2.8E-05; z-score = -0.283) were inhibited. It is noteworthy that mTOR was inhibited at 4 h (Figure 5C) but activated at 1w (Figure 5D). Correspondingly, the expression pattern of myelin protein Plp1, which is a common molecule at both time points, decreased at 4 h but increased at 1 w.

**Figure 4.**
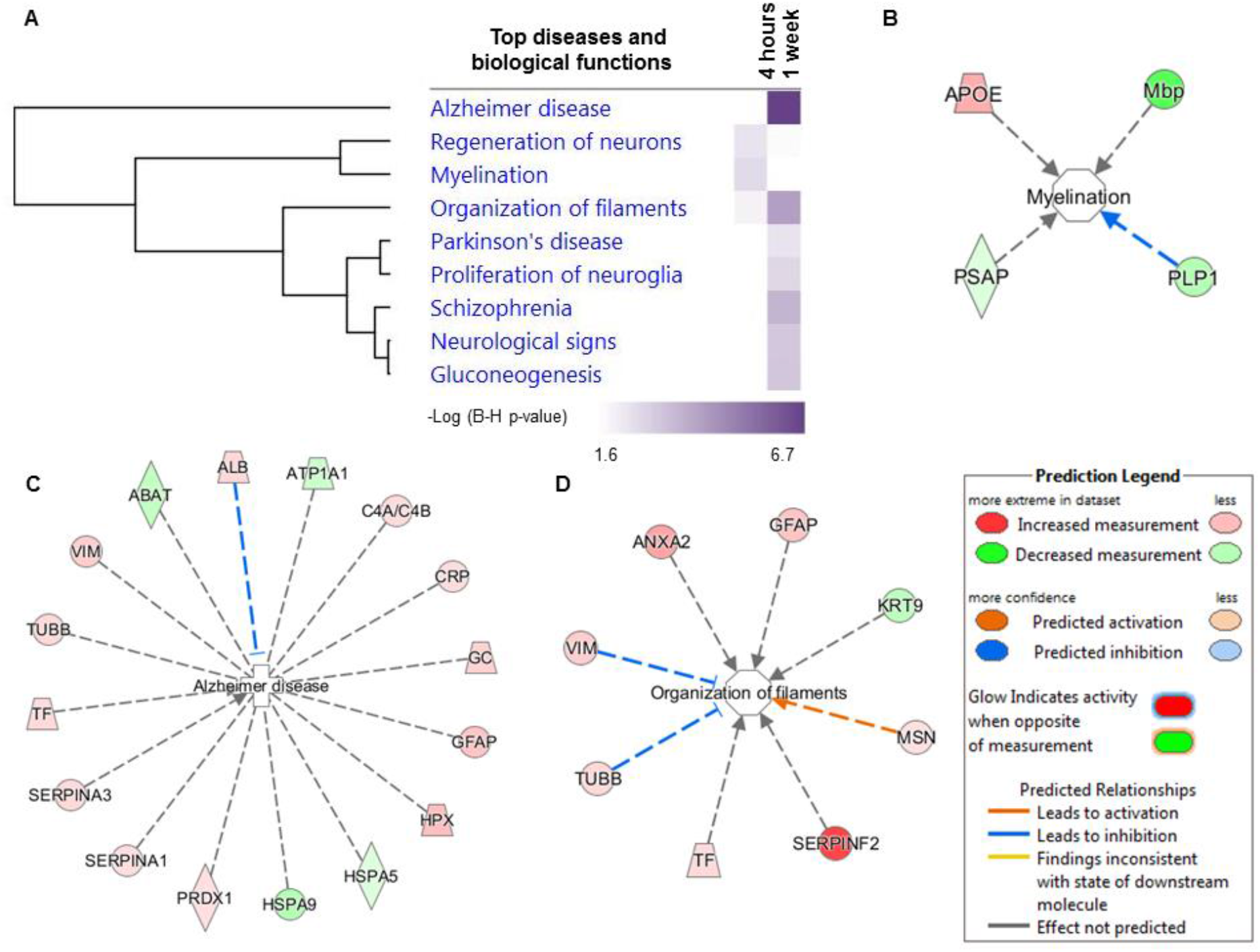
Functional classification of the differentially expressed CSF proteins according to the enriched top diseases and biological functions. (A) Comparison analysis of the significantly affected diseases and biological functions between both time-points following HNK administration. The significance threshold [-log (p-value) >1.6] was determined by IPA analysis. Exemplary interaction networks of top functions at 4h represented by (B) myelination and at 1w represented by (C) Alzheimer’s disease as well as (D) organization of filaments.

**Figure 5.**
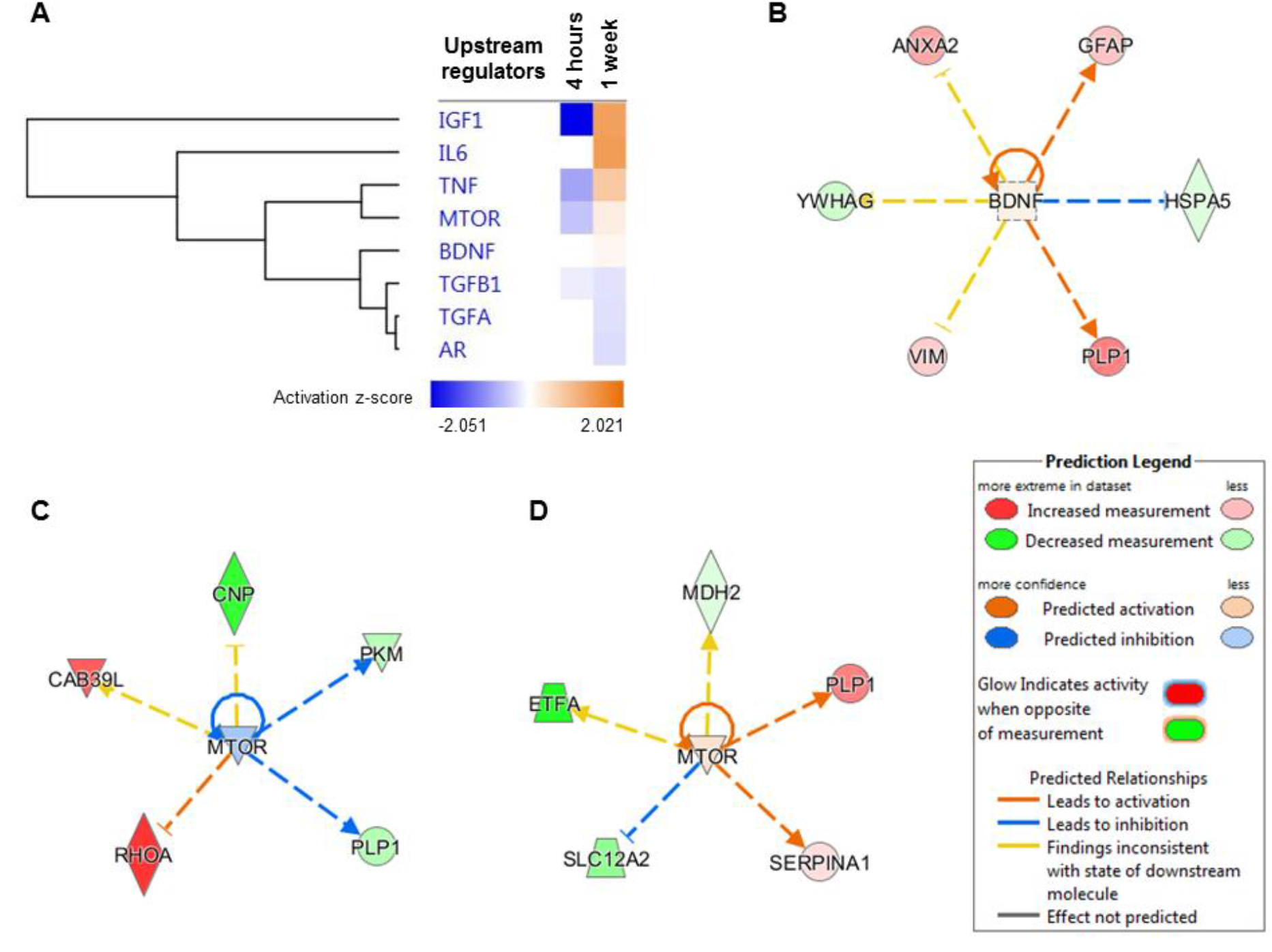
Predicted upstream regulators. (A) Hierarchical clustering of the significantly affected potential upstream regulators observed to experimentally affect the differentially expressed CSF protein clusters at 4h and 1w. The activation z-score of the regulators is depicted as orange (activated) and blue (inhibited). Interaction networks that exemplify the top regulators composed of (B) brain-derived neurotrophic factor (BDNF) and the regulation of mechanistic target of rapamycin (mTOR) at (C) 4h and (D) 1w.

## Discussion

The present study provides first evidence that the DBA/2J mouse strain is a suitable animal model to assess the antidepressant-like effects of rapid-acting antidepressants such as ketamine and HNK. In a next step, we successfully used this model to decode the molecular mechanisms underlying HNK’s rapid acting properties: we characterized the signature and longitudinal changes in the CSF proteome of conscious mice following systemic injection of HNK, which is the most promising metabolite of ketamine. Mechanistically, we identified GR signaling as a putative novel candidate mechanism through which both acute and sustained antidepressant-like effects of HNK might be mediated, thus revealing an intriguing functional link between HNK’s antidepressant mechanism of action and targeted regulation of the stress hormone system.

### Acute and sustained antidepressant-like properties of HNK: importance of experimental model system

In 2016, Zanos *et al.* first described the antidepressant-like properties of HNK (13), which was then further investigated and developed as a putative rapid-acting antidepressant compound with a more favorable side-effect profile than ketamine (11, 12). However, while antidepressant-like effects of ketamine in rodents are relatively robust, the behavioral effects of HNK are more inconsistent and seem to depend on different parameters within the experimental conditions and the mouse strain used (24). Most of the recent studies on ketamine and HNK applied experimental depression models, such as stress or chronic inflammation (16). For example, Chou *et al.* used a learned helplessness paradigm to induce a depressive-like phenotype in rats and demonstrated that HNK decreased depression-like behavior independent of gender (11). Another study using a lipopolysaccharide model of depression was not able to detect any antidepressant-like effects of HNK that were comparable to those of ketamine in male C57/BL6J mice (25). One plausible explanation for those inconsistencies of behavioral effects of HNK in rodents might be the use of different strains and experimental disease models and approaches to mimic clinically relevant conditions.

Adding to this, we could show for the first time, that a single dosage of HNK is equally effective as ketamine in inducing both acute and sustained antidepressant-like effects in DBA/2J mice. Saline-injected animals (i.e. vehicle controls) showed a depression-like behavioral phenotype, which could be attenuated by administration of ketamine and HNK. The clear antidepressant-like effect of both compounds is probably due to the innate high anxiety-like behavior of this inbred mouse strain, which has proven its particular value in psychopharmacological research into antidepressant mechanisms of action (18). Therefore, it is surprising to see that for the rapid-acting compound ketamine and its metabolite HNK, many studies published so far used the C57/BL6 strain, which, under baseline conditions, does not show a typical depression-like behavioral phenotype. Our data support that the DBA/2J strain is a suitable mouse model for studies into the neurobiology of rapid-acting antidepressant compounds. The majority of the published studies revealed antidepressant-like effects of ketamine 24 hours after injection (16). In addition, several studies have shown a sustained, lasting effect of ketamine up to two weeks (26). This prompted us to analyze two different time points and we could indeed show that ketamine and HNK produced both acute (4 hours) and sustained (1 week) antidepressant-like effect in male DBA/2J mice in the FST. Furthermore, ketamine administration significantly impaired locomotor activity, comparable to sedation and other related side-effects that can be prominently seen in patients (10). HNK, in contrast, did not show an impact on locomotor activity in DBA/2J mice, supporting the fact that due to its favorable side-effects profile, HNK might be an ideal candidate to be further tested as rapid-acting antidepressant in clinical use. This prompted us to exclusively focus our in-depth proteomic analysis on HNK.

### Temporal CSF proteome profiles associated with antidepressant-like effects of HNK point to involvement of GR signaling as a putative mechanism of action

As CSF is an accessible biological fluid in both mice and humans that is circulating in close vicinity to the brain, this translational approach was intended to shed light on the molecular pathways underlying HNK’s rapid and sustained antidepressant-like properties. Our results demonstrated that HNK induced time-dependent changes in the murine CSF proteome compared to vehicle-treated control mice.

Our data point, for the first time, to an important role of the GR signaling pathway in the rapid and sustained antidepressant-like properties of HNK. There is convincing evidence in the literature, both from human studies and experimental models, that depression is linked to a dysfunction of the hypothalamus-pituitary-adrenal (HPA) axis, finally resulting in increased levels of circulating glucocorticoids (27). In particular, impaired GR signaling was discussed to be a key mechanism in the pathogenesis of depression (28), which is in line with the hypothesis that antidepressants reduce depressive symptoms by restoring HPA axis function (27). For example, a recent study by Arango-Lievano *et al.* showed that the disruption of GR signaling in rats induced depression-like symptoms and that GR signaling was involved in modulating antidepressant-induced neuronal plasticity needed for reversal of depression-like symptoms (29). While the close link between HPA axis activity and mechanism of action is well established for “classical” antidepressants such as tricyclic antidepressants or serotonin reuptake inhibitors, data on a putative involvement of the stress hormone system in mediating antidepressant-like effects of prototypical rapid acting compounds is sparse. For ketamine, it was described that in addition to its antidepressant-like properties, it was able to restore HPA axis function and to normalize peripheral CORT levels in a mouse chronic stress model (30). So far, data on a putative impact of HNK on modulating GR signaling was lacking. In addition to the proteomic profiling data pointing to an involvement of GR signalling, we here provide a first mechanistic insight into HNK’s effect on GR function: we could show, for the first time, that GR responsiveness *in vitro* was reduced following HNK treatment. Interestingly, a large majority of the proteins implicated in the GR signaling are composed of keratins and these structural proteins were found to be down-regulated following HNK treatment at both time points. Although keratins are traditionally associated with the skin (31), their role in stress signaling has opened a new avenue in neurological disorders. This is supported by recent findings that provided compelling experimental evidence on the role of a specific keratin in CSF as a potential biomarker for Alzheimer’s disease (32). Furthermore, a study by Helenius et al. has lent support to the notion that the expression levels of different keratin subtypes are proportionate to stress in intestinal diseases (33). Apart from keratins, heat shock proteins (HSPs) were also found to be similarly differentially expressed in stress conditions (33). Correspondingly, our results have demonstrated that HSPA5 and HSPA9 were also downregulated in addition to the keratins at the 1w time point. Therefore, based on these results, it is tempting to suggest that downregulation of keratins and also HSPs in our mouse model reflect the HNK-mediated amelioration of stress and this effect lasted for as long as one week. Collectively, these results highlight the crucial effect of HNK in regulating the GR signaling in maintaining the homeostatic balance in the CSF proteome. Among the other differentially expressed candidate proteins, we found Dynll2 which plays a pivotal role in controlling cytoskeletal structures (34) to be upregulated 4 hours after HNK administration.

Several other cytoskeletal proteins like Arpc4, Vim, Tubb5, Klkb1, and Msn were also shown to be induced one week after HNK administration. These findings are in line with recent reports indicating the relevance of cytoskeletal dynamics in translating stressful environmental experiences into persistent changes of molecular and cellular function (35, 36). The fact that we identified a considerable number of cytoskeletal proteins to be specifically regulated by HNK (i.e. putative targets of HNK) could point to their involvement in mediating rapid antidepressant-like effects, which is an intriguing hypothesis considering their crucial role in fine-tuning of synaptic plasticity and function (37). Moreover, we found proteins Mbp, Plp1, and Psap to be altered by HNK administration, with downregulation of Mbp and Plp1 at 4 hours following HNK administration, and upregulation one week later. The latter are proteins playing important roles in myelination and myelin integrity (38). In fact, Weckmann and colleagues provided evidence that ketamine, the parent compound of HNK, increased Mbp protein levels in mice (39). In addition, the antidepressant-like effects of ketamine are known to take effect *via* enhancement of synaptogenesis: a single injection of ketamine triggered mammalian target of rapamycin (mTOR) pathway, which led to antidepressant-like effects by enhancing synaptogenesis in the medial prefrontal cortex of rodents (40-42). The bidirectional amino acid transport with SLC transporter proteins is known to regulate mTOR activation (43). Indeed, we found that mTOR was one of the top predicted upstream regulators, which was activated one week after HNK administration. Correspondingly, transporter proteins Slc25a4 and SLC25A5 were also shown to be upregulated one week after HNK administration. Collectively, these results indicate that the activation of the mTOR pathway might play a pivotal mechanistic role in mediating HNK’s rapid behavioral effects, comparable to what has been described for ketamine.

A limitation of this study is that pooled CSF samples were used for the MS-based proteomics analyses. Although it has been shown that mouse CSF samples can be analyzed individually (44), we pooled the samples to minimize inter-individual variation.

In conclusion, our study substantially advances our knowledge about the molecular and cellular mechanisms through which HNK as a prototypical rapid acting antidepressant compound exerts its acute and sustained antidepressant-like effects. We could confirm the DBA/2J inbred mouse strain to be a suitable animal model for investigations into the neurobiological mechanisms underlying rapid-acting antidepressants. Our translationally relevant approach provides unprecedentedly detailed information on the temporal profile of CSF proteome changes in response to HNK. Finally, by a combination of the hypothesis-free mass spectrometry approach with candidate-driven *in vitro* experiments, we identified, for the first time, GR signaling to be a specific target of HNK. We are confident that those data can be a valuable resource for future studies investigating improved, rapid acting strategies to treat depression.

## Methods

### Animals

Male, adult (8 weeks) DBA/2J mice were purchased from Charles River, France. After arrival at our animal facility, we single-housed all the mice (temperature = 22±2 °C, relative humidity = 50±5 %) and allowed them to habituate to the new environment for at least one week prior to any experiments. We applied a standard 12h dark/light cycle (8am - 8pm light, 8pm - 8am dark). Mice had *ad libitum* access to water and food. All experiments were conducted in accordance with European animal welfare laws and approved by the local animal welfare authority (Landesuntersuchungsamt Rheinland-Pfalz, Koblenz, Germany).

### Drug administration

Murine body weight (BW) was measured 24 h before drug administration. Mice received ketamine hydrochloride (Inresa Arzneimittel GmbH, Germany; 30 mg/kg BW, 5 mg/ml), (2*R*,6*R*)-hydroxynorketaminehydrochloride (Tocris, Germany; 10 mg/kg BW, 2.5 mg/ml), or saline control (0.9% NaCl, Braun, Germany) intraperitoneally.

### Behavioral assessment

#### Forced Swim Test (FST)

Mice were handled for 2 minutes daily for three consecutive days. We then injected them once with either ketamine or HNK or saline. We performed the FST to assess depressive-like behavior 4 h and 1 week after injection. Mice were introduced into a 2-L glass beaker (diameter 13 cm, height 24 cm) filled with tap water (21±1 °C) to a height of 15 cm. We videotaped the mice for 5 minutes and floating, swimming, and struggling behavior was scored by an experienced, observer blinded towards the treatment.

#### Open Field Test (OF)

In a new and independent experiment, mice were handled for 2 minutes daily for three consecutive days. We then injected them once with either ketamine or HNK or saline and immediately placed them into the OF arena (45x45x41 cm). We performed the OF to assess locomotor activity. Mice could move freely and without disturbance for 30 minutes, during which they were videotaped from above. Distanced moved was automatically scored with the tracking software Observer XT12 (Noldus, The Netherlands).

### Blood plasma extraction

We withdrew blood *via* tail nicks into EDTA-coated tubes, as previously described (45). We centrifuged all tubes for 10 min at 10 000 *g* at 4 °C and stored the plasma at -80 °C until further use.

### Corticosterone (CORT) plasma concentration

We measured CORT plasma levels using the Corticosterone ELISA Kit (cat.no. ADI-900-0979; Enzo Life Sciences, USA) according to the manufacturer's protocol in duplicates.

### Statistics of behavioral experiments and neuroendocrine measurements

Data was plotted and descriptive statistics were applied to check for normal distribution of the data (D'Agostino & Pearson omnibus normality test). We analyzed data with a 1- or 2-way ANOVA followed by Bonferroni correction. Alpha was set at 5%, with p values <0.05 considered statistically significant. Data was analyzed using Prism 5 software (GraphPad, USA). Sample sizes are indicated in the figure legends. The values are displayed as mean ± SEM.

### CSF cannula surgery

Ketamine and HNK were equally effective with respect to their antidepressant-like efficacy. Due to the improved side-effect profile of HNK and the gap of knowledge about the neurobiological mechanisms mediating its antidepressant-like activity, we exclusively focused on HNK with respect to the proteome profiling of the CSF.

CSF cannula implantation was conducted in a modified way based on the method by Onaivi and colleagues (19). In brief, mice were anesthetized with Isoflurane (Forene®, AbbVie Deutschland GmbH, Germany) and received the analgesic drug Metacam (Boehringer Ingelheim Vetmedica GmbH, Germany) subcutaneously (0.5 mg/kg BW). The skin was opened and two small holes were drilled into the skull, followed by applying two screws. A third hole was drilled and the intraventricular cannula (internal cannula; Plastics One, USA) was implanted at bregma coordinates 0.0 AP / 0.8 ML / -0.9 VD. Paladur dental cement (Kulzer, Germany) was used to secure the internal cannula to the screws and the skull. The skin lesion was closed and a dummy cannula (Plastics One, USA) closed the internal cannula. Subsequently, the mice received Metacam in the drinking water for one week. After recovery from surgery, mice received new water bottles without the analgesic drug.

### CSF withdrawal and protocol of the main experiment

Mice were handled carefully for 2 minutes per day for 1 week. CSF was withdrawn from all mice and the first sample was discarded to exclude contamination with blood. After three days, mice were randomized into an HNK or saline arm (Supplemental data A). They received an intraperitoneal injection of either HNK or saline and CSF samples were taken 4 hours and 1 week after injection.

CSF withdrawals were conducted as follows: Dummy cannula was removed and tubing (5-10cm in length; Plastics One, USA) with external cannula (additional -1.0 VD (final depth - 1.9 VD); Plastics One, USA) was connected to the internal of the cannula at one end and to a 10μl Model 701 N Hamilton syringe (cat.no. 80365; Hamilton, USA) at the other end. By gently pulling the plunger in steps of 1μl for each 1 minute, CSF was withdrawn. After the plunger reached 10μl, CSF was collected and directly put on dry ice and later stored at -80°C until further analyses. CSF-containing tubes were visually inspected for any blood contamination and excluded if necessary.

### Proteomics sample preparation

The obtained CSF samples of the designated groups were pooled with three biological replicates each, as described in supplemental data A. Briefly, the CSF samples in each assigned biological replicate were pooled equally to a total of 40 μl per replicate. The designated samples were then subjected to one-dimensional gel electrophoresis (1DE) employing precast NuPAGE 4-12% Bis-Tris 10-well mini protein gels (Invitrogen, Germany) with 2-[N-morpholino]ethanesulfonic acid (MOPS) running buffer under reducing conditions at a constant voltage of 150V in 4 °C. Pre-stained protein standard, SeeBlue Plus 2 (Invitrogen, Germany), was used as a molecular mass marker and the gels were stained with Colloidal Blue Staining Kit (Invitrogen, Germany), as per manufacturer’s instructions. The bands in each stained gel were sliced into 14 gel slices per lane with the aim to reduce the complexity and masking effect of the abundant proteins during the MS analysis (Supplemental figure 2). Protein bands were excised (14 bands per replicate) Subsequently, all the excised protein bands were destained, reduced and alkylated prior to in-gel trypsin digestion employing sequence grade-modified trypsin (Promega, USA), as described in detail elsewhere (46-48). Peptides extracted from trypsin digestion were purified with ZipTip C18 pipette tips (Millipore, Billerica, MA, USA) according to the manufacturer’s instructions. The resulting combined peptide eluate was concentrated to dryness in a centrifugal vacuum evaporator (SpeedVac) and dissolved in 10μl of 0.1% trifluoroacetic acid (TFA) for liquid chromatography-electrospray ionization-tandem mass spectrometry (LC-ESI-MS/MS) analysis.

### Discovery proteomics strategy

Peptide fractionation was conducted in the LC system, which consisted of a Rheos Allegro pump (Thermo Scientific, USA) and an HTS PAL autosampler (CTC Analytics AG, Switzerland) equipped with a BioBasic C18, 30 x 0.5 mm precolumn (Thermo Scientific, USA) connected to a BioBasic C18, 150 x 0.5 mm analytical column (Thermo Scientific, USA). Solvent A consisted of LC-MS grade water with 0.1 *%* (v/v) formic acid and solvent B was LC-MS grade acetonitrile with 0.1 % (v/v) formic acid. The gradient was run for 60 min per sample as follows: 0-35 min: 15-40 % B, 35-40 min: 40-60 % B, 40-45 min: 60-90 % B, 45-50 min: 90 % B, 50-53 min: 90-10 % B: 53-60 min: 10 % B.

The continuum MS data were obtained on an ESI-LTQ Orbitrap XL-MS system (Thermo Scientific, Bremen, Germany). The general parameters of the instrument were set as follows: positive ion electrospray ionization mode, a spray voltage of 2.15 KV and a heated capillary temperature of 220 °C. Data was acquired in an automatic dependent mode whereby, there was automatic acquisition switching between Orbitrap-MS and LTQ MS/MS. The Orbitrap resolution was 30000 at *m/z* 400 with survey full scan MS spectra ranging from an m/z of 300 to 1600. Target automatic gain control (AGC) was set at 1.0 x 10^6^ ion. Internal recalibration employed polydimethlycyclosiloxane (PCM) at m/z 445.120025 ions in real time and the lock mass option was enabled in MS mode (49). Tandem data was obtained by selecting top five most intense precursor ions and subjected them for further fragmentation by collision-induced dissociation (CID). The normalized collision energy (NCE) was set to 35 % with activation time of 30 ms with repeat count of three and dynamic exclusion duration of 600 s.

The resulting fragmented ions were recorded in the LTQ. The acquired continuum MS spectra were analyzed by MaxQuant computational proteomics platform version 1.6.3.3 and its built-in Andromeda search engine for peptide and protein identification (50-54). The tandem MS spectra were searched against UniProt *Homo sapiens* (Date, 17 Oct 2018; 20 410 proteins listed) and *Mus musculus* (Date, 17 Oct 2018; 17 001 proteins listed) databases, using standard settings with peptide mass tolerance of ± 30 ppm, fragment mass tolerance of ±0.5 Da, with ≥ 6 amino acid residues and only “unique plus razor peptides” that belong to a protein were chosen (51). Both the *Mus musculus* and *Homo sapiens* databases were utilized with the aim to maximize the protein identification due to limited availability of annotated mouse proteins in a specific database (47). A target-decoy-based false discovery rate (FDR) for peptide and protein identification was set to 0.01. Carbamidomethylation of cysteine was set as a fixed modification, while protein N-terminal acetylation and oxidation of methionine were defined as variable modifications, enzyme: trypsin and maximum number of missed cleavages: 2. The generated MaxQuant output data table “proteingroups.txt” was filtered for reverse hits prior to statistical analysis and, subsequent functional annotation and pathway analyses. The summary of MaxQuant parameters employed in the current analyses is tabulated in Supplemental data B.

### Bioinformatics and functional annotation and pathways analyses

The output of the generated “proteingroups.txt” data from the MaxQuant analysis was utilized for subsequent statistical analysis with Perseus software (version1.6.1.3). First, all raw intensities were log2-transformed and the data were filtered with minimum of three valid values in at least one group and the missing values were imputated by replacing from normal distribution (width: 0.3; down shift: 1.8)), enabling statistical analysis (54). For statistical evaluation, a two-sided Student’s t-test was used for the group comparison with p-values < 0.05 to identify the significantly differentially abundant proteins. Unsupervised hierarchical clustering analysis of the differentially expressed proteins was conducted with log2 fold ratio of the protein intensities according to Euclidean distance (linkage = average; preprocess with k-means) and elucidated in a heat map. The overlaps of the statistically differentially expressed proteins between the databases utilized were manually curated based on the highest significance values. The list of the identified proteins was tabulated in Excel and their gene names were used for subsequent functional annotation and pathways analyses employing Ingenuity Pathway Analysis (v01-04, IPA; Ingenuity QIAGEN Redwood City, CA) (https://www.qiagenbioinformatics.com/products/ingenuity-pathway-analysis) (55). IPA analyses unraveled the protein-protein interaction (PPI) networks, identified the significantly affected canonical pathways, top disease and functions, and predicted upstream regulators associated with the proteins identified to be differentially expressed. Top canonical pathways of the differentially expressed proteins were presented with p-value calculated using Benjamini-Hochberg (B-H) multiple testing correction (-log B-H > 1.3) and a cut-off of ≥ 3 affected proteins per pathway. In PPI networks, proteins molecules are represented by their corresponding gene names and, only PPIs that were experimentally observed and had direct and indirect interactions were used.

### *In vitro* GR activity and gene reporter assays

HT-22 mouse hippocampal neuronal cell line was used as an in vitro model to analyze the molecular effects of HNK on GR transcriptional activity and GR sensitivity to glucocorticoids. Experiments were conducted as described before (56, 57). Briefly, cells were cultivated, transfected with MMTV-Luc and Gaussia-Luc expression constructs using Lipofectamine 2000 (Thermofisher) and treated with HNK in concentrations of 0.1 μM, 1.0 μM, 10.0 μM, and 30.0 μM. Two conditions using culture medium (DMEM, Gibco) supplemented with either standard fetal bovine serum as or with charcoal stripped serum as steroid-free condition were examined to observe HNK effects on the GR-driven transcriptional activity. For evaluating the GR response to hormonal activation, GR-agonist dexamethasone (0.1 nM, 0.3 nM, 1 nM, 3 nM, 10 nM, 30 nM, 100 nM) was added and cells were treated with either HNK (5μM) or vehicle. Data were analyzed using 2-way ANOVA followed by Bonferroni correction. Alpha was set at 5%, with p <0.05 considered statistically significant. Data was analyzed using Prism 5 software (GraphPad, USA).

## Author Contributions

DH, NP, MM designed the research. DH, GT, JN, JE, TJ, AH, and MvdK performed animal experiments. NP, CM, MR performed proteomics experiments. NG performed in vitro experiments. DH, NP, CM, and NG analyzed the data. All authors made a substantial contribution to the interpretation of the data and to the drafting of the manuscript. DH and NP shared first authorship and are listed in alphabetical order.

## Acknowledgements

We are thankful for the technical support by Kathrin Kuna (Mainz, Germany). A portion of the work described herein was carried out by Jens Nadig and Milena Rossmanith in partial fulfilment of the requirements for a medical doctoral degree at the Johannes Gutenberg University Medical Center Mainz, Germany.

DH is supported by the Mainz Research School of Translational Biomedicine (TransMed) with a MD-PhD fellowship. CM is supported by the German Research Foundation (DFG), grant number MA 8006/1-1. TJ is supported by the Focus Program of Translational Neurosciences (FTN) in Mainz with a PhD fellowship. GT is supported by a 2014 NARSAD Young Investigator Grant from the Brain & Behavior Research Foundation and the Danish Council for Independent Research, grant number DFF-5053-00103. MM and KL are supported by the German Research Foundation (DFG) within the Collaborative Research Center 1193 (CRC1193, https://crc1193.de/) and by the Boehringer Ingelheim Foundation.

